# Genomic and ecologic characteristics of the airway microbial-mucosal complex

**DOI:** 10.1101/2022.09.08.507073

**Authors:** Leah Cuthbertson, Ulrike Löber, Jonathan S. Ish-Horowicz, Claire N. McBrien, Colin Churchward, Jeremy C. Parker, Michael T. Olanipekun, Conor Burke, Orla O’Carroll, John Faul, Gwyneth A. Davies, Keir E. Lewis, Julian M. Hopkin, Joy Creaser-Thomas, Robin Goshal, Kian Fan Chung, Stefan Piatek, Saffron A.G. Willis-Owen, Theda U. P. Bartolomaeus, Till Birkner, Sarah Dwyer, Nitin Kumar, Elena M. Turek, A. William Musk, Jenni Hui, Michael Hunter, Alan James, Marc-Emmanuel Dumas, Sarah Filippi, Michael J. Cox, Trevor D. Lawley, Sofia K. Forslund, Miriam F. Moffatt, William O.C. Cookson

## Abstract

Lung diseases due to infection and dysbiosis affect hundreds of millions of people world-wide^1-4^. Microbial communities at the airway mucosal barrier are conserved and highly ordered^5^, reflecting symbiosis and co-evolution with human host factors^6^. Freed of selection to digest nutrients for the host, the airway microbiome underpins cognate management of mucosal immunity and pathogen resistance. We show here the results of the first systematic culture and whole-genome sequencing of the principal airway bacterial species, identifying abundant novel organisms within the genera *Streptococcus, Pauljensenia, Neisseria* and *Gemella*. Bacterial genomes were enriched for genes encoding antimicrobial synthesis, adhesion and biofilm formation, immune modulation, iron utilisation, nitrous oxide (NO) metabolism and sphingolipid signalling. RNA-targeting CRISPR elements in some taxa suggest the potential to prevent or treat specific viral infections. Homologues of human *RO60* present in *Neisseria* spp. provide a possible respiratory primer for autoimmunity in systemic lupus erythematosus (SLE) and Sjögren syndrome. We interpret the structure and biogeography of airway microbial communities from clinical surveys in the context of whole-genome content, identifying features of airway dysbiosis that may presage breakdown of homeostasis during acute attacks of asthma and chronic obstructive pulmonary disease (COPD). We match the gene content of isolates to human transcripts and metabolites expressed late in airway epithelial differentiation, identifying pathways that can sustain host interactions with the microbiota. Our results provide a systematic basis for decrypting interactions between commensals, pathogens, and mucosal immunity in lung diseases of global significance.

## Introduction

### Respiratory infection and immunity

The mucosal surfaces of the airways and lung are extensive and constantly challenged by inhaled microorganisms^7-9^. Overt respiratory infections are the leading cause of death in developing countries, resulting in 4 million lost lives annually^1^. Asthma and COPD each affect more than 300 million people worldwide and are driven by respiratory infections^10^. Two-thirds of individuals exposed to COVID-19 in their home^11^ and half of subjects directly challenged with COVID-19^12^ do not develop infections because of unknown factors.

Upper and lower airways contain a characteristic microbiome^13^ that acts as a gatekeeper to respiratory health^14^. The commensal microbiota regulate immunity in the respiratory mucosa through multiple mechanisms^15-17^. These appear within the first days of life and coincide with susceptibility or resistance to colonisation and infection^18^.

The nose, oropharynx and the intra-thoracic airways form a contiguous tract. The nasopharyngeal mucosa differs histologically and functionally from lower sites^19^, as does its resident microbiota^20^. Pulmonary diseases arise in the intrathoracic airways, whose commensal microbiota are similar to those of the oropharynx^13,21,22^. Up and downward microbial movement occurs between sites^22^. Respiratory pathobionts such *Streptococcus pneumoniae, Haemophilus pneumoniae*, and *Neisseria meningitidis* are commonly carried in the nose and throat without symptoms. The oropharyngeal microbiota do not vary greatly between individuals and are organised into co-abundance networks that may share similar niches^5^. Microbial community dysbiosis with overgrowth of pathobionts has been shown in asthma, COPD and other pulmonary disorders^14,23^.

Airway commensal organisms have not previously been systematically cultured or sequenced, limiting the structured study of interactions between bacteria, viruses, fungi and mucosal immunity in clinical samples or in model systems. In this paper we describe such systematic exploration, substantially extending what is known about core constituents of airway microbiomes. Our study design is summarised in Supplementary Figure 1. We have used mucin-enriched media to culture and sequence novel taxa that together account for 75% of the abundance of airway commensal organisms. Functional characterization, evolutionary analyses and comparison with amplicon sequencing in representative human samples extends the scope of these results.

## Results

### Culture collection and isolate novelty

Lower airway bacteria were cultivated from bronchoscopic brushings from two asthmatics and three healthy individuals from the Celtic Fire Study (described below). We used a limited range of media with and without 0.5 % mucin, followed by incubation in standard atmosphere or an anaerobic workstation to capture 706 isolates. Those without overlapping 16S rRNA gene sequences were transferred to the Wellcome Sanger Institute and whole genome sequenced with assembly using Bactopia (v 1.4.11).

Out of 256 cultures with successful whole-genome sequencing, five appeared mixed and were removed. After removing duplicates on a threshold of 99.5% nucleotide identity 126 unique strains remained. Forty-four isolates were annotated to species level in accordance with MIGA^24^ (TypeMat and NCBIProk) and with GTDBtk. A further 30 species were identified by either MIGA (TypeMat and NCBIProk) or GTDBtk. All isolates were assigned to genera in the TypeMat or NCBI prokaryotes database with p<0.05. Among these samples we classified 49 *Streptococcus*, ten *Veillonella*, nine each of *Gemella* and *Rothia*, eight *Prevotella*, six each of *Neisseria, Micrococcus* and *Pauljensenia*, five each of *Haemophilus* and *Staphylococcus*, three *Granulicatella*, two each of *Actinomyces, Cutibarterium* and *Fusobacterium* and one *Cuprividis, Leptotrichia, Microbacterium* and *Niallia*, respectively (Figure 1a).

**FIGURE 1.**
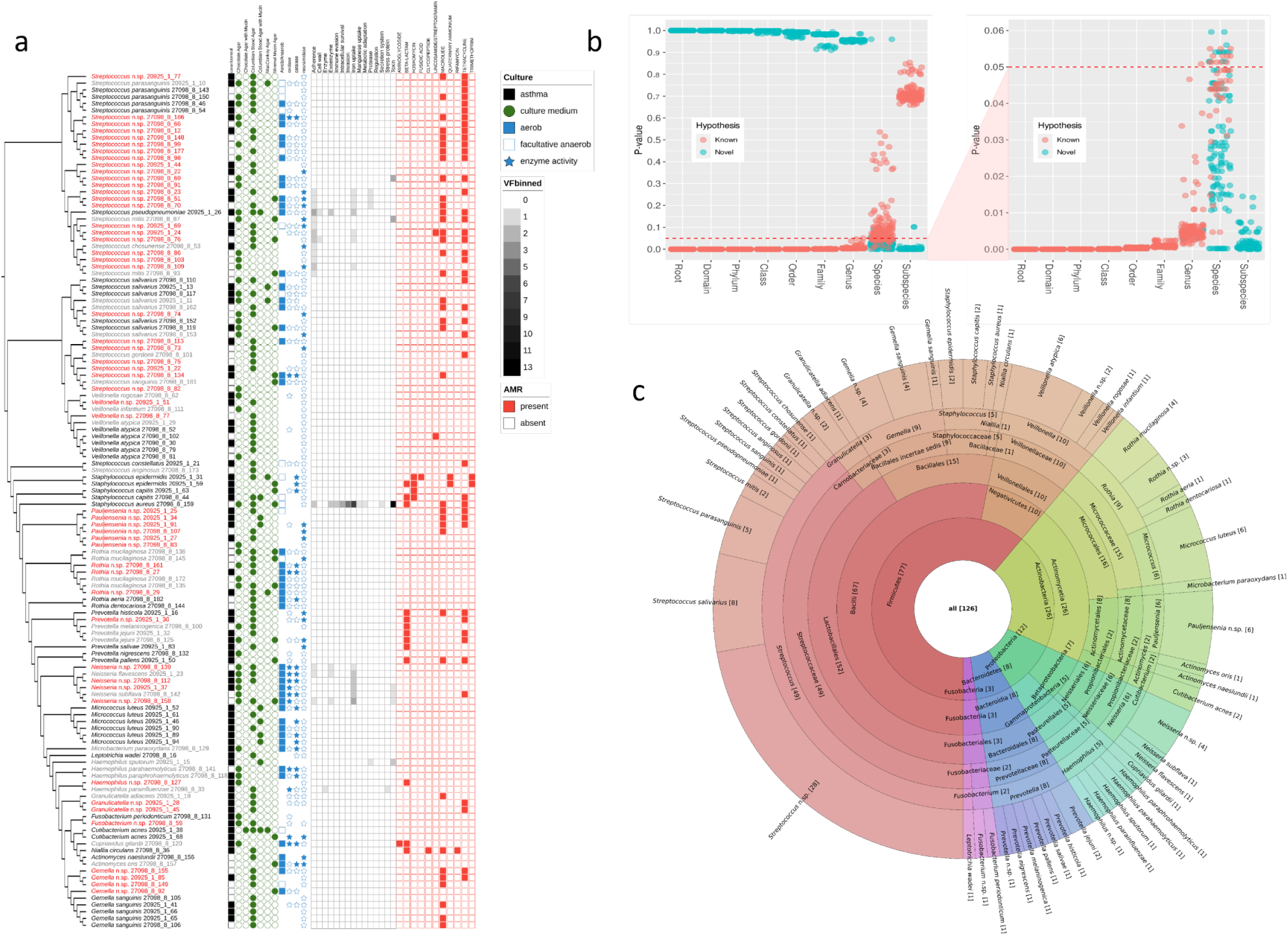
Genomic characteristics of airway mucosal bacteria a) Culture collection phylogeny based on average nucleotide identities between genomes with 1000bp fragment length. Putatively novel species are highlighted in red (indicating that it is not related to any species in the TypeMat DB or NCBI Prok DB (p<0.05) when assessed using MIGA and not assigned to a known species or incongruent species assignment using gtdbtk). Greyed-out isolates are not fully supported by MIGA and gtdbtk. Genome completeness and contamination are displayed as a barchart. AMR finder was used to identify antimicrobial resistance genes at the protein level (red panel). Virulence factors were identified using the VFDB and ariba databases and binned into 15 categories (heatmap). Asthma status of the host is indicated in the black asthma/control panel. Cultivation conditions are indicated in green circles for selected growth media, blue rectangles for aerobic and white rectangles for anaerobic cultivation. Positive gram staining for GNB, GNC, GPB, GPC and other gram staining is shown in black circles. Neuraminidase activity was tested if a blue star is present and is filled for positive test and white for negative test. b) Taxonomic novelty as calculated by MIGA using TypeMat reference. The scatterplot shows support (P-value, vertical axis) for each taxon relative to complementary hypotheses that this taxon is a previously known one (red markers) or a novel one (cyan markers) at each taxonomic level (horizontal axis). Many of the isolate collection constitute novel species within known genera. c) Composition of bacteria isolated and cultivated from five subjects. Counts are shown for all lineages from species level (outer circle) to phylum level (inner circle) in squared brackets. The ETE3 toolkit was used to fetch taxonomic lineages for all genera of cultured isolates ^72^. The number of unique species was summed up and visualised along with their lineages using Krona tools^73^.

Fifty-two isolates could not be assigned with p<0.05 to known species in the reference databases^24^ (Figure 1b). Twenty-eight of the putative novel species were contained within the *Streptococcus* genus, six within *Pauljensenia* (not previously recognised to be prevalent in the airways), and four each within *Neisseria* and *Gemella* (Figure 1c).

Comparison of the full sequences of our streptococcal isolates with 2477 public *Streptococcus* spp. sequences showed that the organisms were widely distributed amongst *S. infantis*, S. *oralis, S. mitis, S. pseudopneumoniae, S. sanguinis, S. parasanguinis*, and *S. salivarius* (Supplementary Figure 2).

### Isolate characteristics

#### Kegg ontology of isolate genomes

We used the eggNOG (evolutionary genealogy of genes: Non-supervised Orthologous Groups) mapper tool (as previously for large-scale systematic genome annotations^25^) to assign by transfer 5,531 Kegg Ontology (KO) annotations for the 126 isolates. We encoded these in a binary matrix indicating presence or absence (Supplementary Table 1) and constructed an isolate phylogeny after removing 254 zero-variance KOs either present or absent in all isolates and reducing identical KO presence/absence to single examples before hierarchical clustering with the Manhattan distance metric and complete linkage. The Dynamic Tree Cut algorithm^26^ identified 15 clusters of isolates that recovered known phylogenetic relationships (Figure 2a). Based on the observed 16S rRNA gene sequence similarity, we further divided one *Streptococcus* cluster into two (Step I and Strep II, Figure 2a). Relative KO enrichment was estimated for each of the 16 clusters by contingency table analysis.

**FIGURE 2.**
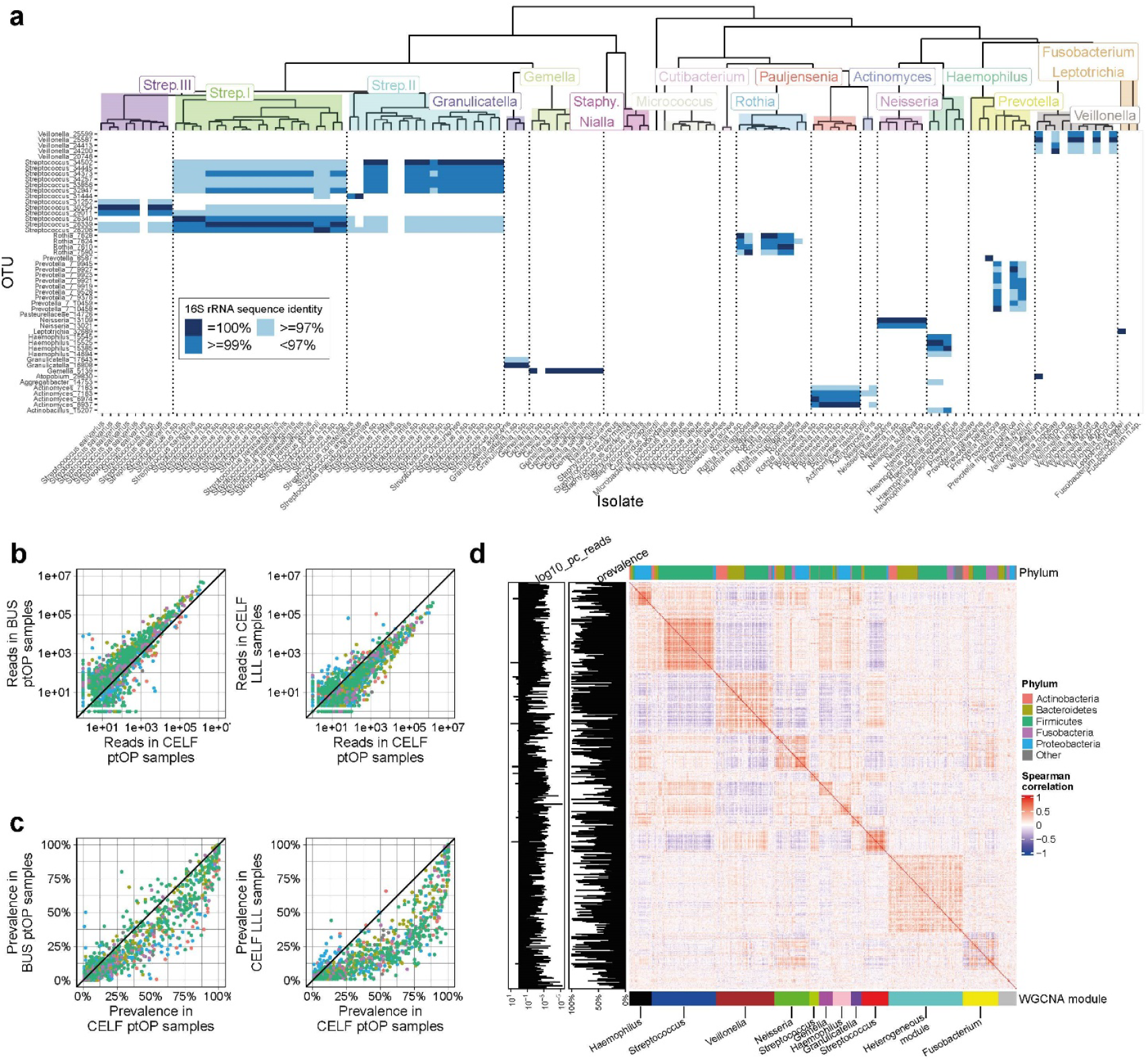
Ecology and structure of airway microbial communities **a**) Mapping of the 50 most abundant OTUs onto 126 novel airway isolates. Isolates are grouped into 15 clusters according to distance and branching order of their inferred Kegg Ontology (KO) gene content. OTU/isolate nt identity is shown as 95-97% (light blue), 97-99% (medium blue) and 100% (dark blue). The complex relationship between OTUs and isolates reflects multiple copies of the 16S rRNA gene in different taxa, but in general captures KO phylogenetic structures. **b**) Comparison of abundance (left) and prevalence right) of bacterial OTUs in populations from northern (CELF) and southern (BUS) hemispheres. The species distribution is similar between the CELF and BUS studies **c)** comparison of abundance (left) and prevalence right) of bacterial OTUs in the posterior oropharynx (ptOP) and the left lower lobe (LLL) in CELF subjects. The relative abundance of organisms in ptOP is very similar to those in the LLL, although absolute abundance is an order of magnitude lower in the LLL. Lower abundance OTUs in the CELF dataset are more prevalent in the upper than lower airways **d)** Spearman correlations between the abundance of organisms in the CELF ptOP samples, showing a high degree of positive and negative relationships between OTUs that is the basis of WGCNA network analysis. Common phyla are colour coded at the top of the matrix, and WGCNA modules (named for the most abundant membership) are at the bottom. Network module membership may be dominated by a single phylum (e.g., the *Haemophilus* or *Streptococcus* modules) or contain mixed phyla (e.g., the *Veillonella* module).

Annotation for the 5,277 informative KOs (including duplicates removed during clustering) (Supplementary Table 2a) identified 247 uncharacterised proteins (Supplementary Table 2b). Features of particular interest among the known genes are summarised below.

#### Biofilms

Biofilm formation is a feature of respiratory pathogens, archetypically *Pseudomonas* spp. in patients with cystic fibrosis. Biofilm-associated genes were also common in the commensal collection (Supplementary File 2b). Ninety genes were annotated with “biofilm” in their KO pathway descriptions, with *cysE* (serine O-acetyltransferase), *vpsU* (tyrosine-protein phosphatase), *luxS* (S-ribosylhomocysteine lyase), *trpE* (anthranilate synthase component I) and *PYG* (glycogen phosphorylase) present in >75%% of isolates. Amongst the most abundant organisms, *Haemophilus* and *Prevotella spp*. had distinctive profiles of other biofilm pathway genes (Supplementary Table 2b).

#### Antimicrobial resistance and virulence

Many of our isolates contained known genes for antimicrobial resistance (AMR) against tetracyclines and macrolides. *Staphylococcus, Prevotella* and *Haemophilus* spp. also possessed beta-lactam resistance (Figure 1a). Virulence factors and toxins were concentrated in *Streptococcus, Staphylococcus, Haemophilus* and *Neisseria* spp. (Figure 1a). Although these annotations neither guarantee the genes in question are expressed nor that they drive clinically relevant AMR or virulence, they do indicate such potential.

#### Antibiotic and toxin synthesis

Competition between bacteria is fundamental to maintaining stable communities^27^. Genes with a KO pathway annotation for antibiotic synthesis (n=33) were present in many genera (Supplementary Table 2c). Arachin biosynthetic genes included *acpP* (acyl carrier protein) which was present in 120 isolates and *auaG* in 7 (mostly *Staphylococcus* spp); *rifB* (rifamycin polyketide synthase) present in 20 (*Veillonella* and *Staphylococcus* spp.); *BacF* (bacilysin biosynthesis transaminase) present in 12 (*Staphylococcus* and *Gemella* spp.); and *sgcE5* (enediyne biosynthesis protein E5) present in in 12, mostly *Haemophilus* spp.. Bacteriocin exporter genes *blpB* and *blpA* were present in 35 and 31 isolates respectively, predominately *Streptococcus* and *Pauljensenia* spp. (Supplementary Table 2d).

Toxins and antitoxin genes were common in the collection (Supplementary Table 2d), without distinctive enrichment in particular genera. They included homologues of antitoxin *YefM* (57 isolates); exfoliative toxin A/B *eta*, (57 isolates); toxin *YoeB* (51isolates); antitoxins *HigA-1* (31) and *HigA* (30); antitoxin *PezA* (26); toxin *RtxA* (15); antitoxin *HipB* (14); toxin *YxiD* (13); antitoxin *<u>CptB</u>* (12); antitoxin *Phd* (11); and toxin *FitB* (10). These have not been previously recognised in commensal organisms and differ from the toxin spectrum of known airway pathogens^28^. They may have significant influences on the mucosa as well as other organisms.

#### Nitric Oxide

Nitric oxide (NO) is a central host signalling molecule in the airways, where it mediates bronchodilation, vasodilation, and ciliary beating^29^. NO exhibits cytostatic or cytocidal activity against many pathogenic microorganisms^30^ and NO elevation in exhaled breath is used as a clinical marker for lower airway inflammation. Many isolate genes encoded NO reductases (Supplementary Table 2f), including norB (27 isolates); norV (11), norQ (5), norC (1) and norR (1). The hmp gene, encoding a NO dioxygenase, was present in 39 organisms. These enzymes may mitigate the antimicrobial activities of NO or affect host bronchodilation and mucus flow.

#### Iron and heme

Iron is an essential nutrient for humans and many microbes and is a catalyst for respiration and DNA replication^31^. Host regulation of iron distribution through many mechanisms serves as an innate immune mechanism against invading pathogens (nutritional immunity)^31^.

We identified 47 genes with “iron” in their KO name (Supplementary Table 2f). Those found in >75% of isolates were *afuC* (iron (III) transport system ATP-binding protein), *ABC*.*FEV*.*P* (iron complex transport system permease protein), *ABC*.*FEV*.*S* (substrate-binding protein), and *ABC*.*FEV*.*A* (ATP-binding protein). A further 19 genes were identified as members of “heme” pathways (Supplementary Table 2g).

*Haemophilus* spp. require heme for aerobic growth and possess multiple mechanisms to obtain this essential nutrient. These genes may play essential roles in *Haemophilus influenzae* virulence^32^. In our isolate collection *sitC* and *sitD* (manganese/iron transport system permease proteins) and *fieF* (a ferrous-iron efflux pump) were only found in *Haemophilus* spp., as were *ccmA, ccmB, ccmC, ccmD* (heme exporter proteins A, B, C and D) and *hutZ* (heme oxygenase). These are potential therapeutic targets.

#### Sphingolipids

The sphingolipids constitute an important class of bioactive lipids, including ceramide and sphingosine-1-phosphate (S1P). Ceramide is a hub in sphingolipid metabolism, and mediates growth inhibition, apoptosis, differentiation and senescence. S1P is a key regulator of cell motility and proliferation^33^.

Sphingolipids play significant roles in host antiviral responses^34,35^ and resistance to intracellular bacteria^36^. Their importance in humans is exemplified by a major childhood asthma susceptibility locus that upregulates *ORMDL3* expression^37^. ORMDL3 protein acts as a rate limiting step in sphingolipid synthesis^38^ and the *ORMDL3* locus greatly increases the risk of HRV-induced acute asthma^39^.

*De novo* synthesis of sphingolipids is recognised in human bowel bacteria ^40^ and maintains intestinal homeostasis and microbial symbiosis^41^. In the skin, commensal *S. epidermidis* sphingomyelinase makes a crucial contribution to skin barrier homeostasis^42^. Based on KO annotations, we did not find obvious SPT homologues in our isolates but identified 12 genes with putative roles in sphingolipid metabolism (Supplementary Table 2g). Of these, *SPHK* (sphingosine kinase, present in 12 isolates) which metabolises sphingosine to produce S1P; and *ASAH2* (neutral ceramidase, present in 7 isolates) have potential roles in modifying host inflammation and repair. These may interact with the *ORMDL3* disease risk alleles described above.

#### Immune inhibition

Several genes present in the isolates may directly affect host immunity. These were enriched in *Prevotella* spp. (Supplementary Table 2h) and included immune inhibitor A (*ina*), a neutral metalloprotease secreted to degrade antibacterial proteins; *Spa* (immunoglobulin G-binding protein A), *sbi* (immunoglobulin G-binding protein Sbi); *omp31* (outer membrane immunogenic protein); *blpL* (immunity protein cagA); and *impA* (immunomodulating metalloprotease).

A conserved commensal antigen, β-hexosaminidase (HEXA_B), has a major role in induction of anti-inflammatory intestinal T lymphocytes^43^, and is present in 59 of our isolates with enrichment in *Prevotella, Streptococcus* and *Pauljensenia* spp.

#### Autoantigens

Systemic lupus erythematosus (SLE) and Sjögren syndrome are chronic autoimmune inflammatory disorders with multiorgan effects. Lung involvement during the course of the disease is frequent^44^. Our *Neisseria* isolates contain a 60 kDa SS-A/Ro ribonucleoprotein (Supplementary Table 2a) that is an ortholog to the human *RO60* gene, a frequent target of the autoimmune response in patients with SLE and Sjögren’s syndrome.

Other bacterial genomes contain potential Ro orthologs^45^, and a bacterial origin of SLE autoimmunity has been suggested^46^. Here, the abundance of *Neisseria* spp. in human airways and their close proximity to the mucosa are of interest, as is a recent report that the lung microbiome regulates brain autoimmunity^47^, and an earlier observation that T cells become licensed in the lung to enter the central nervous system^48^.

It is relevant that products of cognate microbial-immune interactions in the airways have direct access to the general arterial circulation through the left side of the heart, whereas molecules and cells arising from the gut undergo extensive filtration and metabolism in the liver before accessing more distant sites.

#### CRISPR genes

Most respiratory viruses, including SARS2-Cov-19, have RNA genomes, and RNA-targeting CRISPR vectors have the potential to prevent or treat viral infections ^49^. Type III RNA-targeting system elements (such as cas10, cas7, csm2 and csm5)^50^ are present in our isolates (particularly *Fusobacteria* and *Prevotella* spp.), as is the Type II system element cas9 (Supplementary Table 2i).

### Isolates in the context of airway communities

#### Community coverage

We sought context to our culture collection within the ecological variation of different geographic and anatomical locations. We studied airway microbial communities in 66 asthmatics and 44 normal subjects recruited from centres in Dublin (48 subjects), Swansea (48 subjects) and London (16 subjects) (collectively known as the Celtic Fire Study (CELF)). Swabs were taken from the posterior oropharynx (ptOPs) and bronchoscopic brushings from the left lower lobe (LLL) in all subjects. When tolerated the left upper lobe (LUL) was also brushed in 52 subjects. We compared the European CELF microbial communities to 527 ptOP samples from an adult population sample in Busselton, West Australia (BUS)^5^. Operational Taxonomic Units (OTUs) were identified by sequencing the 16S rRNA gene amplicon and compared with the assembled genomes from our culture collection.

In the CELF ptOP samples, 17 operational taxonomic units (OTUs) covered >70% of the abundance and 41 OTUs covered >85% (Supplementary Table 3). Coverage was less complete in LLL and LUL samples (respectively 64% and 50% at the 70% threshold), due to the expansion of *H. influenzae* (OTU Haemophilus_14694) and *Tropheryma whipplei* (OTU Glutamicibacter_5653) in the pulmonary samples, particularly those from asthmatics (Supplementary Table 3).

Fifteen of the most abundant 17 OTUs were mapped to at least one isolate using a 99% nt identity, and 11 of the next 24 mapped to a cultured organism. Genera of moderate abundance (2.8%-0.4% of the total) yet to be cultured include *Fusobacterium, Selenomonas, Alloprevotella, Porphyromonas, Leptotrichiaceae, Megasphaera, Lachnospiraceae, Solobacterium*, and *Capnocytophaga*. We have previously shown that *Leptotrichia, Selenomonas, Megasphaera* and *Capnocytophaga* spp. are reduced in abundance in asthmatic ptOP samples^5^. Future isolation is desirable to test if they are indicator species or direct contributors to respiratory health.

OTUs corresponding to isolates for *Staphylococcus, Micrococcus* and *Cupriavidus* spp. had minimal representation in the community OTU analyses, although *S. aureus* is a recognised lung pathogen. Their appearance in the isolates may represent oral or skin contamination or assertive growth in culture.

Mapping of the 50 most abundant OTU sequences onto the 126 isolates revealed complex relationships that reflect multiple copies of the 16S rRNA gene in different taxa^51^ (Figure 2a). In general, however, OTU assignment reflected the principal KO phylogenetic structures, and referencing of OTU communities to our isolate genomes may still inform on community functional capabilities.

The 16S rRNA gene sequences poorly detected the extensive diversity of *Streptococcus* spp. in airways, as noted previously^5^. However, combinations of OTUs can be seen to form “barcodes” (Figure 2a) that may refine *Streptococcus* spp. identification into their three main KO phylogenetic groups.

#### Biogeography and community structure

The taxa defined by OTUs, and their relative abundances were similar in CELF ptOP and CELF LLL samples, and to the normal population in BUS ptOP (Figure 2b and Figure 2c). Other than the most abundant organisms, the prevalence of most OTUs was lower in the LLL than in the ptOP (Figure 2c). The mean bacterial burden was much higher in ptOP samples from CELF than in the LLL (log10 mean 7.86±0.07 vs 5.06±0.05), consistent with previous studies^13,21,22^.

Strong correlations and anti-correlations were present between the abundances of OTUs in data from each site (exemplified for CELF ptOP samples in Figure 2d, and previously shown for the BUS ptOP results^5^). We used WGCNA analysis to find networks (named arbitrarily with colours) within these correlated taxa. Network structures were consistent in the CELF and BUS ptOP communities (Figure 2e and 2g), but less distinct in the lower airway samples (Supplementary Figure 3) where taxa were less diverse and of lower abundance (Supplementary Figure 3).

Networks often contained closely related species but also extended beyond phylogenetically related organisms (Figure 2g). For example, in the CELF ptOP networks (Figure 2b) there are phylogenetically homogeneous modules of *Streptococci* (blue, red and greenyellow), *Gemella* (magenta), *Haemophilus* (black and pink) and *Granulicatella* (purple).

Of interest is the brown module in the CELF ptOP samples, which contain multiple *Prevotella* and *Veillonella* spp. of high abundance. The presence of biofilm elements in *Prevotella* spp. described above supports a hypothesis that these organisms may adhere to form a basic “commensal carpet” of the airways^5^.

Both the CELF ptOP and BUS ptOP networks recovered the phylogenetic relationships found in the KO analysis amongst *Streptococcus* isolates. The three clusters of *Streptococcus* isolates (Strep. I-III) map to distinct sets of OTUs using sequence similarity (Figure 2a), and this similarity is also uncovered in the WGCNA network modules in both ptOP networks (Supplementary Figure 4).

### Dysbiosis

Subtle alterations in bacterial community composition (“dysbiosis”^52^) are recognized in many diseases with microbial components. Community instability and inflammation in the presence of mild viral infections^10^ can be added to the recognized features of loss of diversity and pathobiont expansion in asthma and COPD. We therefore sought novel insights into airway dysbiosis in our subjects from genomic sequencing of the commensal organisms.

We used Dirichlet Multinomial Mixtures (DMM)^53^ to assign airway community components on all samples from the BUS and CELF subjects. Samples formed predominantly into two clusters (Airway Community Type 1 and 2, ACT 1 and 2) (Figure 3a). The main drivers for the two clusters were identified as *Streptococcus, Veillonella, Prevotella* and *Haemophilus* spp. in descending order of relative abundance across all samples. ACT1 was dominated by *Streptococcus, Veillonella* and *Prevotella* in 410 samples; whilst ACT2 was dominated by *Streptococcus, Veillonella* and *Haemophilus* in 478 samples (Figure 3a). Principal coordinates analysis based on Bray-Curtis-distance (β-diversity) of the airway microbiota confirmed significant overall compositional differences between the two community type clusters (PERMANOVA p-value > 0.001) (Figure 3b).

**FIGURE 3.**
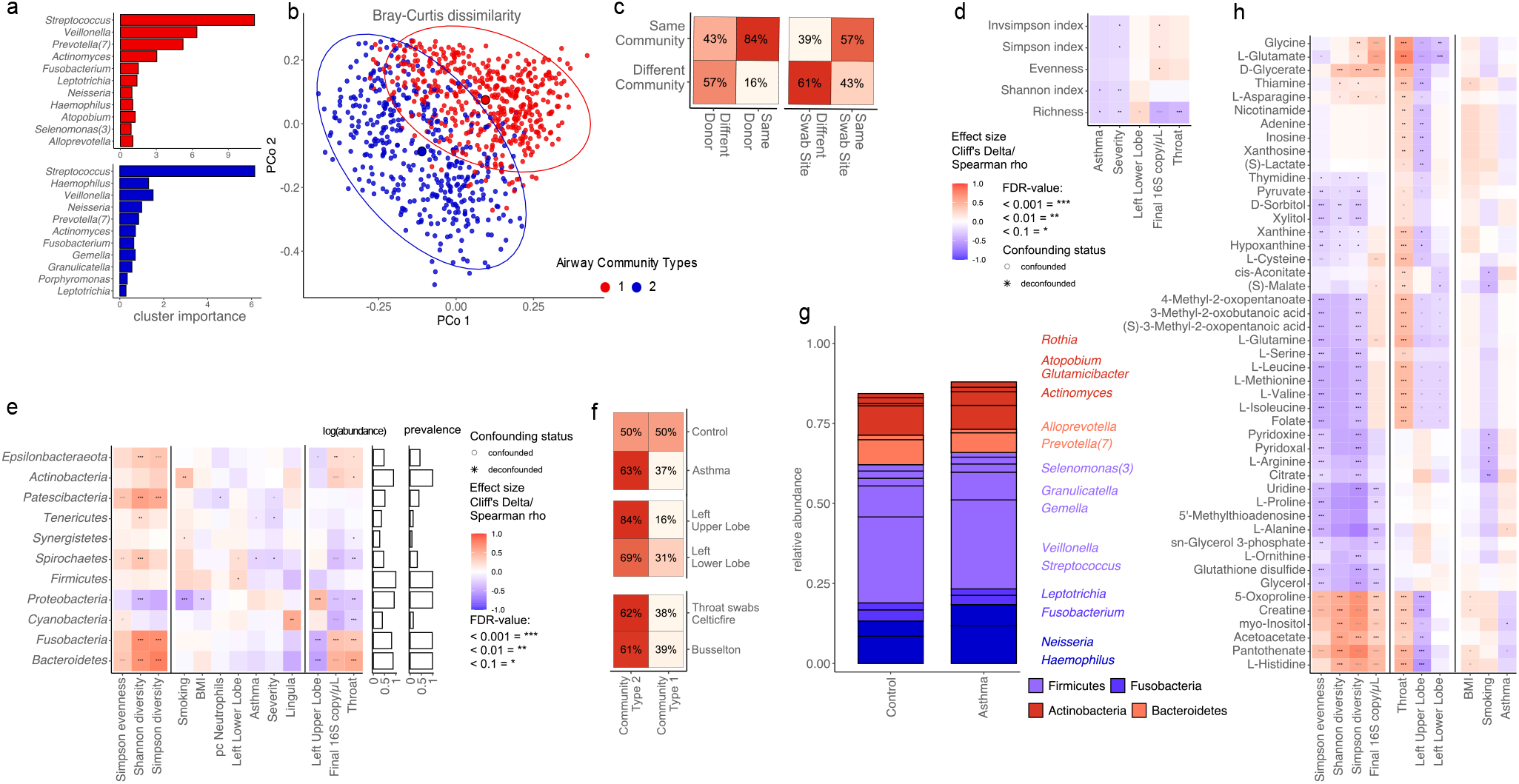
Microbial features of dysbiosis **a)** Main drivers of Dirichlet multinomial model-based airway communities. **b)** beta diversity based on Bray-Curtis dissimilarity principal coordinate analysis showing separation of the two communities. **c)** alpha diversity measures **d)** Consistency of airway community assignment between samples of same and different donor(left) and sampling site (right). **e)** Proportion of community assignments between throat samples of different study origin (left), sampling site (middle), disease group (right). **f)** Univariate associations of CELF 16S samples binned on phylum level to metadata. **g)** relative abundance of most abundant phyla (left) and genera (right) based on CELF samples 16S rRNA. **h)** Univariate metabolite associations based on binning of CELF 16S rRNA sequences onto isolate annotation.

We investigated effects varying between airway sites in the CELF subjects. To assess effects on alpha diversity measurements (Figure 3c) and the relative abundance of specific bacterial taxa (Figure 3d), we conducted univariate analysis to relate evenness and richness (Figure 3c) and phylum level taxon abundance (Figure 3e) to the metadata describing the CELF subjects. Metadata features describing clinical phenotypes and sample origin were often strongly collinear, and so we assessed found associations in turn for retained significance with each potential confounder, using a nested rank-transformed mixed model test previously implemented as a publicly available tool^54^ and considering repeated sampling of patients as a random effect.

Congruence analysis with regards to ACT assignment of CELF samples (Figure 3c, left) confirmed consistency in assignment for samples coming from the same donor (χ^2^ < 0.005) or the same sampling site (χ^2^ < 0.005). We saw pervasive effects both on alpha diversity indices and phylum level of the tested predictors (Figure 3c & 3d). Importantly, the Shannon index and richness were significantly decreased with asthma status and severity (MWU FDR < 0.1) (Figure 3c). We saw an increase (although not significantly) of the *Proteobacteria* Phyla associated with asthma status (Figure 3d), in line with the taxonomic profile of patients with asthma vs. healthy controls (Figure 3e). This is consistent with many reports of *Proteobacteria* excess in asthmatic airways^13,14,55^.

ACT proportions from CELF samples (Figure 3e, right) differed significantly with regards to asthma status (n= 285, χ^2^ < 0.05) and sampling site (n=176, χ^2^: Left lower lobe < 0.1, left upper lobe < 0.001, false discovery rate (FDR) controlled at 10 %).

### Mucosal factors

Next, to relate our charted microbiome diversity to the salient properties of its ecosystem niche, we sought host components of the microbial-mucosal interface by serial measurements of global gene expression and supernatant metabolomics during full human airway epithelial cell (HAEC) differentiation in an air-liquid interface model (ALI). We hypothesised that the transition from monolayer to ciliated epithelium over 28 days would be accompanied by progressive expression of genes and secretion of metabolites for managing the microbiota.

We found 2,553 significantly changing transcripts organised into eight core temporal gene clusters of gene expression (Limma, 3.22.7) (Figure 4a and Supplementary Table 4). Four clusters showed late peaks of expression and three of these (CL2, CL4 and CL5) contained many genes that are likely to interact with the microbiome (Supplementary Table 4). Transcripts in the other upgoing cluster (CL3) were elevated early and late in differentiation and were enriched for genes mediating cell mobility and localisation. Genes of particular interest in the other upgoing clusters are as follows.

**FIGURE 4.**
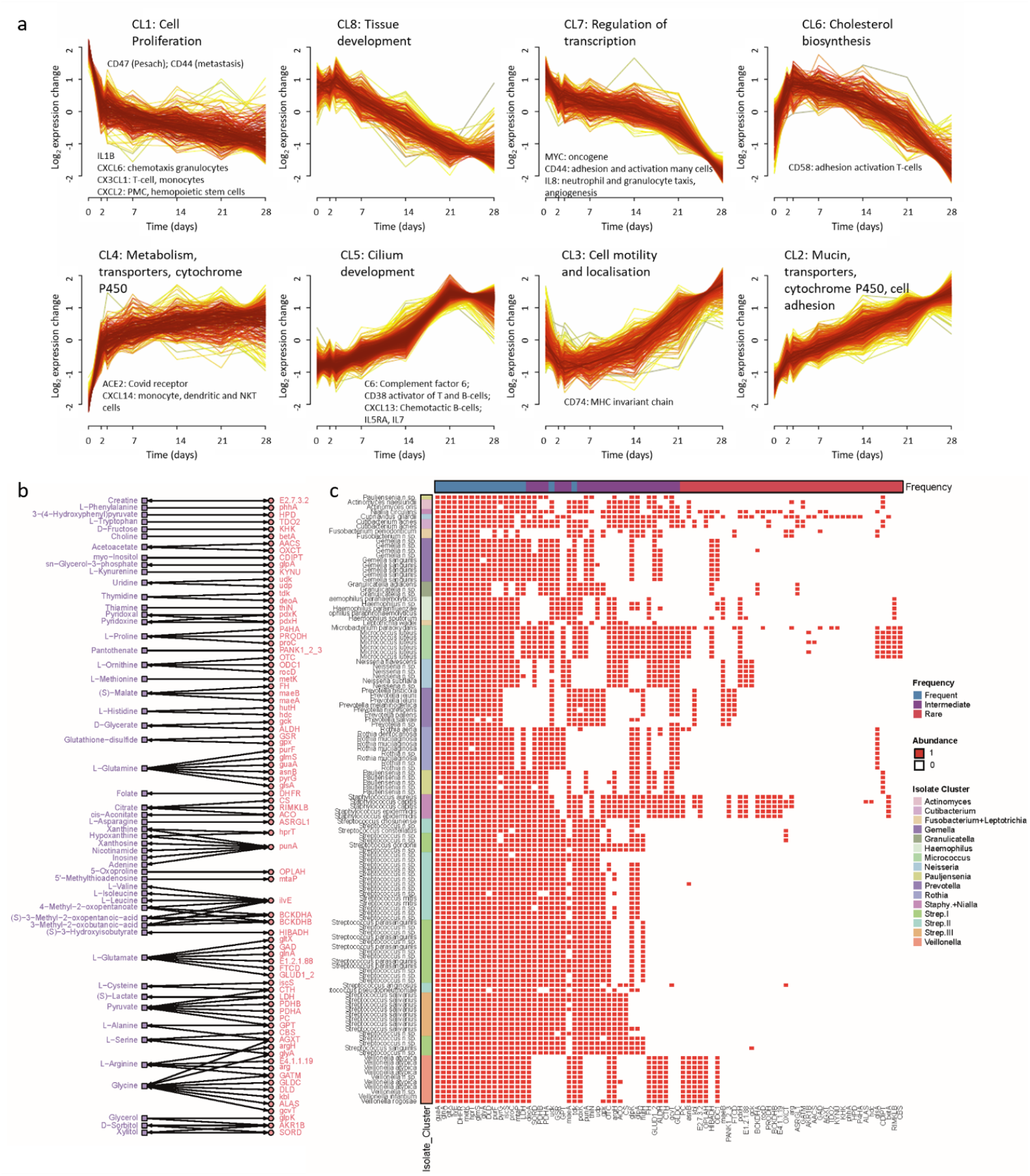
Gene and metabolite abundance during airway epithelial development a) Global gene expression was measured at 7 times over 28 days in an air-liquid model of epithelial differentiation (monolayer to ciliated epithelium). A total of 2,553 transcripts, summarised by 8 core temporal profiles, showed significant variation in abundance during mucociliary development. Hallmark functional roles are shown for each cluster. Clusters CL2, CL3, CL4 and CL5 show late peaks of expression and contain genes that can interact with the microbiome. Upregulated chemokines and immune-function genes are also noted within the clusters. b) Metabolites (square) measured in the supernatant of the fully differentiated airway cells were linked to genes (circle) identified in bacterial isolates. Arrows indicate if the reactions were reversible or irreversible, with metabolites as substrates and products. These networks were built based on KEGG pathways. c) Binary heatmap displaying the presence (1) or absence (0) of genes (columns) identified in the genomic sequences of bacterial isolates (rows). Bacterial isolates are organised into Kegg Ontology phylogeny clusters (see Figure 2). Gene annotations (top) indicate the frequency of the gene: ‘frequent’ for genes in >75% of isolates, ‘intermediate’ for genes in 25-75% of isolates and ‘rare’ for those in <25% of isolates.

#### Mucins and ciliary development

Mucosal mucins are central to mucosal function and integrity, providing a source of nutrients and sites for tethering of commensals^56^, at the same time as restricting the density of organisms through upward flow by beating cilia^57^. Interaction of mucins with <u>microbiota</u> plays an important role in normal function^56^, and direct cross-talk between microbes and mucin production is likely^57^.

In our ALI model, progressive up-regulation of the major secreted respiratory mucins *MUC5AC* and *MUC5B* in CL2 was accompanied by the membrane associated *MUC20* (Table 1, Supplementary Table 4). In contrast, CL5 contained 3 membrane-associated mucins (*MUC13, MUC15, MUC16*). These mucins do not form gels and are anchored to the apical cell surface where they present a glycoarray for selective interactions with the microbial environment^56^.

Within CL5 we also found 17 gene families and 175 genes with putative roles in ciliary function, ciliogenesis, or spermatogenesis (Supplementary Table 4). Mutations in many of these genes are known to cause primary ciliary dyskinesia (PCD)^58^, which results in recurrent pulmonary infections. Other genes in this list are candidates for mutation in cases of PCD without known cause.

#### Immune related genes

The most significant effects (top hits) in CL2 included *ENPP4* (which promotes haemostasis); *ALOX15* (which generates bioactive lipid mediators including eicosanoids); *GLIPR2* (which enhances type-I IFNs); *MPPED2* (a metallophosphoesterase active in infection); *INSR* (insulin receptor); and *MIR223* (an inhibitor of neutrophil extracellular trap (NET) formation in infection) (Table 1, Supplementary Table 4).

Immune-related genes significantly expressed in CL5 included complement factor 6 (*C6*) which forms part of the membrane attack complex. C6 deficiency is associated with *Neisseria* spp. infections. *CD38* was also highly expressed, and its product is an activator of B-cells and T-cells.

#### Detoxification and transportation

Top hits in CL4 include *ADH1C*, an alcohol dehydrogenase; *GSTA2* with a known role in detoxification of electrophilic carcinogens, environmental toxins and products of oxidative stress by conjugation with glutathione; *ACE2*, the SARS2-Cov-19 binding site which cleaves angiotensins; and *PIK3R3* which phosphorylates phosphatidylinositol to affect growth signalling pathways (Table 1, Supplementary Table 4).

CL4 contains five members of the cytochrome P450 families with potential roles in detoxification of microbial products, including *CYP2F1* (which modifies tryptophan toxins and xenobiotics); *CYP4X1* (unknown substrates); *CYP4Z1* (benzyl esters); *CYP4F3* (Leukotriene B4); and *CYP2C18* (sulfaphenazole). Also in CL4 were transporters *SLC10A5* (substrate bile acids); *SLC27A2* (fatty acids); *SLC1A1* (glutamate); *SLC4A11* (borate); *SLC25A4* (ADP/ATP in mitochondria); *SLC45A4* (sucrose); *SLC25A28* (iron); and *SLC39A11* (zinc).

Enrichment of genes for detoxification and transport was also present within CL2, which included *CYP4B1* (substrate fatty acids and alcohols); *CYP4V2* (fatty acids); *CYP2A13* (nitrosamines); *CYP2B6* (xenobiotics); *CYP26A1* (retinoids); and *CYP4F12* (arachidonic acids). Transporters included *SLC40A1* (iron); *SLC13A2* (citrate); *SLC15A2* (small peptides); *SLC12A7* (KCl co-transporter); and *SLC35A5* (nucleoside sugars).

#### Neuronal development

The bronchial mucosa is innervated with unmyelinated fibres that detect airway luminal substances^59^ and mediate smooth muscle tone, mucus secretion, and cough. Stimulation of airway sensory nerve endings also generates the release of proinflammatory molecules^60^ (“neural inflammation^61^”).

A basis for innervation can be seen in top hits from CL2, which included *ENPP5* and *HECW2*, which have putative roles in development of airway sensory nerves (Supplementary Table 4). Interestingly, CL2 and CL4 together contained ten members of the protocadherin beta gene family (*PCDHB2, PCDHB3, PCDHB4, PCDHB5, PCDHB10, PCDHB12* and *PCDHB18P* in CL2; *PCDHB13, PCDHB14*, and *PCDHB15* in CL4). Interactions

between protocadherin beta extracellular domains specify self-avoidance in specific cell to cell neural connections^62^, and their abundant presence here may regulate singular neural-mucosal cell coherence.

### Intersection of mucosal and microbial metabolomic pathways

Metabolites are central to biological signalling, and so we used the same time-series model of AEC differentiation to measure levels of metabolites released into the culture media of the cells (Supplementary Table 5).

We then mapped these ALI culture metabolites to enzymes in matching bacterial pathways identified within the KO of isolate genomes (Figure 4b), based on direct reactions, as substrates or products. Notable interactions include amino acids, nucleotides and compounds involved in energy metabolism. The metabolite-related KOs exhibited distinctive patterns within the isolate phylogeny (Figure 4c).

Enrichment of these KOs onto global human and bacterial KO pathways with iPath^63^ is shown in Supplementary Figures 4a and 4b. These suggest folate biosynthesis to ubiquitous amongst airway organisms, valine, leucine and isoleucine metabolism to be of intermediate importance and alanine, aspartate and glutamate metabolism to be rare functions amongst the isolates.

Extrapolation of metabolic activities was possible from binning 16S abundance onto the isolate KOs using an approach modelled on the PICRUSt program^64^, revealing metabolite profiles that distinguished measures of diversity and location within upper or lower airways (Figure 3i), as well as distinctive features of asthma and dysbiosis.

## Discussion

Our results provide an inventory of the genomic and metabolomic capacities of the respiratory commensal bacteria and of the fully differentiated respiratory epithelium that they inhabit. Known mechanisms through which commensal microbiota regulate immunity include activations of inflammasomes^15^, Nod2 and GM-CSF^16^, and chemokines^17^. Such factors are present in neonates during microbial differentiation with subsequent susceptibility or resistance to infections^18^. Our study suggests multiple other host factors for managing microbial growth, including metabolites.

It is to be expected that other pathways, particularly involving immune signalling, will only become evident when bacteria and the mucosa are grown together. With our representative airway isolate collection, our findings set a stage for systematic investigation of the dynamic interplay within members of the microbial-mucosal complex in health and in the protean respiratory conditions that arise at the border between the environment and the lung.

Metagenomic sequencing has been the cornerstone of many studies of the bowel microbiota, but non-purulent sputa (airway secretions) typically contain <5% microbial DNA^65^ and cellular samples such as brushings and biopsies will contain even less. Abundant pathogens and commensals may nevertheless be identified by sequencing, albeit at great depth ^65,66^. Our results will greatly improve metagenome assembly and allow assays of individual microbial activities through metatranscriptomics.

Microbial community dysbiosis with diversity loss and overgrowth of pathobionts is recognised in asthma, COPD and other pulmonary disorders^14,23^. HRV infections are the major precipitant of acute exacerbations of asthma^67,68^ and of COPD^69,70^ yet have trivial effects in most individuals. Here we have found networks of interacting bacteria that are attenuated in the lower airways, possibly presaging loss of stability^71^. The hypothesis can now be tested that airway dysbiosis and microbial community instability predisposes to catastrophic dysregulation of airway microbiota and inflammatory processes during acute exacerbations of lung disease. Eventually, the successful repair of dysbiotic airway microbial communities may help treat asthma and prevent lung infections.

## Methods

### Microbial culture

After sampling. bronchial brushes for extended culture were immediately placed in 15 ml centrifuge tubes with 2 ml sterile saline solution (0.9% w/v) and immediately transported to the laboratory for processing. Samples were mixed on a vortex mixer twice for 5 seconds. On duplicate plates, 100 µl of the saline was plated on Columbian blood agar (5% horse blood), chocolate agar or minimal agar with 0.5 % (w/v) mucin. One set of plates were incubated at 37 & C in standard atmosphere while the other set was incubated at 37 & C in an anaerobic workstation (Don Whitley DG250). Colonies were selected from 24 hours to 168 hours by appearance, streaked out on their corresponding media and incubated for a minimum of 48 hours. Plates were then colony selected again and Gram stained. Aerobic isolates were tested for oxidase and catalase activity. DNA was extracted from brain heart infusion broth for aerobes and sodium thioglycollate media for the anaerobes. Any isolate which failed to grow in liquid medium were grown on solid medium and an inoculation loop was used to scrape growth off the surface of the agar prior to DNA extraction.

### Whole Genome Sequencing bacterial isolates

Whole genome sequencing was carried out at the Wellcome Sanger Institute, using the HiSeq X platform and generating paired-end read lengths of 151bp. Genomes were *de novo* assembled using Bactopia^74^ (v 1.4.11). Taxonomic classification and quality control were performed using MiGA (http://microbial-genomes.org/) with the TypeMat database. Isolates appearing to contain multiple genomes were discarded.

For all assemblies the average nucleotide identity was computed using fastANI^75^ (v 1.3) with a fragment length of 500bp and clustered on 99.5% average nucleotide identity. For every cluster, sequencing data of every entity (isolate) were pooled and processed using Bactopia (v 1.4.11) with default settings. Taxonomic annotation and novelty scores were computed using MiGA with the TypeMat database as well as the NCBI Prokaryote genome database for comparison. Functional annotation was performed using prokka (v 1.14.6) as implemented in Bactopia; and eggnog-mapper^76^ (v emapper-1.0.3-40-g41a8498) using diamond (v 0.9.24) for the alignments, reducing the search space to the domain of bacteria. Antimicrobial resistances were annotated using amrfinder (v 3.8.4) and ARIBA (v 2.14.5) using the CARD database (v 3.0.8). Virulence factors were computed using the VFdb core dataset (v) and binned into higher functional entities using a custom perl script.

Phylogenetic analysis of the isolates was performed using the Bacsort pipeline (https://github.com/rrwick/Bacsort). First, fastANI distances were computed with a fragment length of 1000 bp and a maximum distance of 0.2. A phylogenetic tree was constructed using as implemented in the R-package ape^77^ (v 5.6-2). The tree was visualized using the Interactive Tree of Life (iTol)^78^. Small ribosomal subunits were extracted from assembled genomes using Metaxa2 and aligned with CELF OTUs using BLAST with 100% percentage nucleotide identity, e-value=1e-10, and length ≥206 bp.

### Kegg Ontology and isolate phylogeny

From the eggnog-mapper output we derived 5,531 Kegg Ontology (KO) annotations for the 126 isolates which we encoded in a binary matrix indicating presence/absence. We removed 254 zero-variance KOs (that were either present in all or no isolates) and performed hierarchical clustering of the isolates with the 5,023 remaining KOs using the Manhattan distance metric and complete linkage. The distance matrix was calculated after removing 2,313 KOs that had identical presence/absence to at least one other isolate. The distance matrix was calculated after removing 2,313 KOs that had identical presence/absence to at least one other isolate. The Dynamic Tree Cut algorithm^26^ identified 15 clusters of isolates that recovered known phylogenetic relationships (Figure 2a). These 15 clusters were then mapped to the OTUs using the 16S rRNA gene sequence similarity (Figure 2a). Based on OTU similarities, one *Streptococcus* cluster was split into two additional clusters, resulting in a final set of 16.

We then identified characteristic KOs that were over-or under-represented in each cluster relative to all other clusters. We scored cluster i and KO j using a 2×2 contingency table, where a: number of isolates in cluster i containing KO j; b: number of isolates in cluster i without KO j; c: number of isolates not in cluster i containing KO j and d: number of isolates not in cluster j without KO j; from which we calculated odds ratios (ORs) using ad/bc. 0.5 was added to cells with zero counts (the Haldane-Anscombe correction). Log10(OR) was used a summary statistic to rank the KOs by importance for a given cluster. The 2,313 duplicate KOs were assigned the same score as their duplicated counterpart used to construct the distance matrix.

### Human study populations

Samples included in this study were collected from two study populations, The microbial pathology of asthma study (Celtic Fire, CELF) and the Busselton health study, a long running epidemiological survey in South-Western Australia (BUS).

The CELF study was a multicentre, cross-sectional study of asthmatic adults and healthy controls. Participants were recruited from 3 UK centres, Connolly Hospital, Dublin; The Royal Brompton Hospital, London; and Swansea University Medical School, Swansea. Ethical approval for the study was granted by the London-Stanmore Research Ethics Committee (reference 14/LO/2063). All subjects provided written informed consent. Subject groups were: healthy subjects (non-smokers and current smokers; asthmatic patients taking short-acting beta agonists only (BTS Step 1) ; asthmatics on moderate dose of inhaled corticosteroid (ICS) (up to 800 µg/day of beclomethasone propionate (BDP equivalent)± long-acting β-agonist LABA (BTS Step 2/3); asthmatics on high dose ICS (ICS dose >=1600 µg/day) + LABA ± other controllers (theophyllines, LTRA, LAMA) (BTS Step 4); and asthmatics on high dose ICS (ICS dose >=1600 µg/day) + LABA ± other controllers + oral prednisolone ± anti-IgE treatment (BTS Step 5). Severe asthma was defined as BTS step 4 or 5. Exclusion criteria were: Asthmatic subjects must be non-smokers or ex-smokers with < 5 pack-years smoking; BMI>35; diagnosis of rheumatoid arthritis, allergic bronchopulmonary aspergillosis, or Churg-Strauss syndrome; drug therapy with beta-blockers, ACE inhibitors, anti-asthma immune modulators other than steroids; antibiotics within 4 weeks of study; acute exacerbations of asthma within past 4 weeks; history of an upper or lower respiratory infection (including common cold) within 4 weeks of baseline assessments; confounding occupations (such as baking); and significant vocal cord disorder.

Participants were invited to initial assessments prior to bronchoscopy. A posterior oro-pharyngeal (ptOP) swab was taken from each participant immediately before the bronchoscopy commenced. During bronchoscopy, two bronchial brushings were taken from the left lower lobe (LLL) of each subject. If tolerated, two further brushes were taken from the left upper lobe (LUL). An additional bronchial brush from the left lower lobe of five study participants from The Royal Brompton Hospital were processed for extended bacterial culture (described below).

All other samples were stored at -80°C within 1 hour of collection. Those harvested at The Royal Brompton Hospital were transported stored directly to the Asmarley Centre for Genomic Medicine (ACGM) at the same site. Samples at other sites were stored locally at -80°C for a maximum of 6 months prior to transport to the ACGM on dry ice.

Investigation of the BUS subjects was as previously described^5^. ptOP swabs were collected with the same protocols as CELF from 527 individuals. After local storage at -80°C, ptOP swabs were transported on dry ice to the ACGM for further processing.

### DNA extraction and quantification

Microbial DNA extraction from Celtic Fire samples was carried out using a hexadecyltrimethylammonium bromide (CTAB) and bead-beating double extraction using phase lock tubes. Bacterial isolates were extracted using a single extraction method. Full details of extraction protocols for each sample type are outlined in Cuthbertson *et al* 2020 (Protocols.io). Bussleton throat swabs were extracted using the MPBio DNA extraction kit for Soil, as previously described^5^. DNA was stored at -20°C until processing. Microbial DNA quantification was carried out using a SYBR green 16S rRNA gene qPCR^79^.

### Microbial 16S rRNA analyses

16S rRNA gene sequencing was performed on the Illumina MiSeq platform using dual barcode fusion primers and the V2 500 cycle sequencing kit. Sequencing was performed for the V4 region of the 16S rRNA gene as previously described^5,79^. Sampling and extraction controls, PCR negatives and mock communities were included on all sequencing runs.

All samples and controls from both the Celtic Fire and BUS datasets were included in this analysis and were processed through the QIIME 2.0 analysis pipeline.

Sequences were quality trimmed to 200bp using trim-galore (Version 0.6.4) and joined with a maximum of 10% mismatch and a minimum of 150 base pair overlap using joined_paired_ends.py (Version 1.9.1). Data was quality checked using FASTX Toolkit (Version 0.0.14) prior to de-multiplexing.

Reads were dereplicated and open reference OTU clustering was performed in QIIME 2. Chimeric sequences were identified and removed, leaving borderline calls in the analysis. Phylogeny was aligned using mafft followed by consensus taxonomic classification. The Biom file, tre file and taxa identifications were exported for further analysis.

Processed data was transferred to R (Version 3.6.3) and uploaded into Phyloseq (Version 1.3). Reads unassigned or assigned to Archaea at the kingdom level were removed before further analysis along with reads identified as Chloroplast or Mitochondria. All OTUs with less than 20 reads (reads present in less than <2% of the samples (n = 1,174)) were removed from further analysis.

Contaminant OTUs were identified using Spearman’s correlation between bacterial biomass with number of reads per samples. OTUs were considered to be contaminants with a Benjamini-Hochberg corrected P-value of <0.05 and a correlation value of >0.2.

Due to the nature of the differences in the extraction and sequencing protocols between BUS and CELF studies, contaminants were investigated in the whole dataset and in CELF and BUS separately. OTUs identified using the individual datasets were removed from further analysis (Table S2). The “Prevalence” method in Decontam (Version 1.6) with a threshold of 0.1 and controlling for study, identified a further 55 OTUs contaminant OTUs associated with negative controls. All OTUs identified were checked and found to be consistent with contamination^80^.

### Community analyses of 16S rRNA sequences

OTU counts were rarefied to the size of the smallest retained sample (discarding samples with too few reads) to obtain the relative abundances of the microbiota in each sample accounting for read depths.

Univariate analysis was done using metadeconfoundR (https://github.com/TillBirkner/metadeconfoundR), relative abundances were tested for univariate associations with clinical variables, requiring Benjamini-Hochberg adjusted FDR < 0.1 and the absence of any clear confounders. Only major taxa and OTUs detected after rarefaction in at least 10% of samples were used.

Within metadeconfoundR, as described elsewhere^54^ non-parametric tests were used for all association tests as the data was not normally distributed. For discrete predictors, the Mann-Whitney test (two-categorical variables) or the Kruskal-Wallis analysis of variance (more than two categorical variables) were used. For pairs of continuous variables, a non-parametric Spearman correlation test was used. Benjamini-Hochberg False Discovery Rate control (FDR) was applied to control for multiple testing controlling the family-wise error rate at 10%.

Hierarchical clustering on the relative abundance profiles were used to establish grouping patterns of the different study samples, including an updated adaptation of the approach used to define “enterotypes” in the human gut, this so called pulmotyping was performed using the Dirichlet Multinominal package, fitting a Dirichlet-multinomial model on the count matrix of genus relative abundance to classify genus abundance based on probability. Each count x in the matrix corresponds to a feature (of n features in total) in the composition observed in the replicate sample. Replicates are grouped into k groups. This parameterization of the Dirichlet distribution for k parameters corresponds to the expected proportions of each of the features (e.g., a particular taxon) in group k, and is an intensity that is shared among all features. The hyperprior for the k parameters at the ‘topmost’, or most inclusive, level of the model hierarchy is another Dirichlet distribution with equal prior probability for each feature within the composition. These distributions together form a hierarchical model for relative abundances among samples used to cluster all samples into different pulmotypes. The chi-square test implemented in base R was used to test for significant differences in the resulting pulmotype distribution between samples grouped by disease status.

Redundancy-reduced isolate abundance/sample (from 16S) and annotation isolate to KEGG KOs were used to generate a sample to KO projection. The projection was mapped to KOs involved in generating the metabolites highlighted by the ALI experiments^64^, by multiplying taxon abundances with the KO presence/absence matrix to yield functional potentials and a proxy for expected metabolite turnover. MetadeconfoundR analysis of this matrix was then carried out together with clinical metadata accompanying the OTU abundance analysis.

### Airway epithelial cell culture

Primary normal human bronchial epithelial (NHBE) cells (Promocell, Germany) derived from a 26-year old adult were grown on collagen coated flasks using the Airway Epithelial Cell Growth Medium Kit (Promocell, Germany) supplemented with bovine pituitary extract (0.004ml/ml), epidermal growth factor (10 ng/ml), insulin (recombinant human) (5 μg/ml), hydrocortisone (0.5 μg/ml), epinephrine (0.5 μg/ml), triiodo-L-thyronine (6.7 ng/ml), transferrin, holo (human) (10 μg/ml) & retinoic acid (0.1 ng/ml) (Promocell, Germany) and Primocin (Invivogen, France).

At passage 3, NHBE cells were seeded onto 12 mm Transwell inserts with 0.4 µm pore polyester membranes at a density of 2.5×10^5^ cells/insert. Cells were maintained in ALI medium, a 50:50 mixture of ALI x2 media (Airway Epithelial Cell Basal Medium with 2 supplement packs added (without triiodo-L-thyronine and retinoic acid supplements) and 1 ml BSA (3 µg/ml)) and DMEM supplemented with retinoic acid (15 ng/ml) (Sigma Aldrich, Gillingham, UK). Cells were fed apically and basolaterally until 100% confluent, after which they were fed exclusively basolaterally with apical media removed. This was defined as ‘Day 0’, the start of the ALI culture. Media was changed three times a week for 28 days, at which stage full differentiation had occurred. At seven points during culture we performed transepithelial electrical resistance (TEER) measurements, took apical washings for ELISA measuring MUC5AC, harvested triplicate wells for gene expression microarray analysis and qPCR for MUC5AC mRNA as well as harvested quadruplicate wells and culture supernatants for metabolomics analysis. NHBE cell pellets and 200ul basolateral supernatants were snap-frozen in liquid nitrogen and stored at -80°C for metabolomic analysis.

All cell culture experiments were regularly tested for mycoplasma contamination using LOOKOUT® Mycoplasma PCR Detection Kit (Sigma-Aldrich, USA) for mesothelioma cell culture and PCR Mycoplasma Test Kit I/C (Promokine, Germany) for NHBE cell culture.

### Metabolomics analysis

Metabolic profiling performed by Metabolon Inc (NC, USA) followed their standard protocols. NHBE cell samples were analysed using LC-MS and GC-MS methods. All samples were given unique identifiers and bar-coded for tracking throughout the analysis pipeline. The Metabolon LIMS system was used to extract raw data, identify peaks and process QCs. Metabolites were identified by comparing retention times, *m/z* and chromatographic data to library entries of purified standards and recurrent unknown entities. All library matches were confirmed with interpretation software and the assigned compounds were curated. Metabolite data from cell lines were normalised by cell density and missing values, below the limit of detection, were imputed with the lowest detected value for the corresponding variables for subsequent analysis.

Analyses were performed using R (version 4.1.1). The MetaboSignal package^81^ was utilised to link media metabolites to KOs via their shortest paths, according to KEGG pathways. These pathways were filtered to display only direct reversible and irreversible reactions. Metabolites and KOs were mapped to human and microbial metabolic pathways using iPath 3.0 (https://pathways.embl.de/)^63^.

### Transcriptomics of NHBE

Approximately 200ng total RNA (with the exception of one sample in which 100ng total RNA was used) was prepared for whole transcriptome microarray analysis using the Ambion WT Expression kit. Purified cRNA yield was assessed using an Agilent 2100 Bioanalyzer and then taken forward for reverse transcription to yield sense-strand cDNA. A total of 5.5µg of sense-strand cDNA was fragmented and labelled using the Affymetrix GeneChip WT Terminal Labelling Kit prior to hybridization to the GeneChip ST2.1 Array. Micorarray libraries were hybridised, washed, stained and imaged using the Affymetrix Genetitan.

Analyses were carried out in R (version 3.1.0). Raw data was imported into R and quality control carried out using arrayQualityMetrics (version 3.20.0), detecting outlier arrays that are likely to skew data upon normalisation. Any outlier arrays were excluded and the corresponding samples re-processed and run on arrays until all samples had successfully passed quality control. QC-passed arrays were normalised by Robust Multichip Average (RMA) using Affymetrix Power Tools (version 1.12.0). Probe-sets that had below-median levels of expression in all arrays were removed. Differential expression was determined using linear modelling of the time-course using the Limma package (version 3.20.0)^82^. All *P* values are corrected for multiple testing; using a method derived from Benjamini and Hochberg’s method to control the false discovery rate^83^.

Transcripts were clustered based on their expression patterns over the time-course using a soft-clustering approach (MFUZZ)^84^. Gene ontology was determined by the HOMER (Hypergeometric Optimization of Motif EnRichment, version 4.7) program^85^. Fold-change per gene ontology term was determined by: (number of target genes in term / total number of target genes) / (total number of genes in term / total number of genes in background list).

Temporal variation in gene expression was assessed by fitting a temporal trend using a regression spline with 3 df (Limma, 3.22.7). P-values were adjusted for multiple testing, controlling the false discovery rate (FDR) below 1%. TC annotations were compiled from NetAffx (access date 30/06/2020) and hugene21sttranscriptcluster.db (8.5.0). Common temporal expression patterns were sought amongst differentially expressed genes using the unsupervised classification technique Mfuzz (2.26.0), informed by the minimum distance between cluster centroids (Dmin).

### Network Analysis

Co-abundance networks were constructed using Weighted correlation network analysis (WGCNA)^86^. We constructed WGCNA co-abundance networks separately using the CELF ptOP, CELF LLL and BUS ptOP samples, including any OTUs that appeared in 20% of samples in at least one of these four subsets (646 OTUs). Spearman correlation was used to construct the WGCNA adjacency matrices. OTU reads were transformed using log(x+1) prior to WGCNA analysis.

## Author Contributions

MFM and WOC planned the overall study structures; TDL suggested building a culture and sequence collection of airway bacteria, and led sequencing at the Wellcome Sanger Centre; LC, CC, MC and MFM designed and carried out microbial culture of airway samples; CC has catalogued and biobanked the organisms; SKF led bioinformatic strategy for microbial sequences, which were carried out by UL, JI-H, ThB and TiB with advice from SaF; MTO carried out analyses of metabolomic data, with guidance by MD; CMcB carried out the microbial community analyses from the Celtic Fire Study with input from CC, JI-H and LC; CB, OO’C, JF, GD, KL, JC-T, JMH, RG, and FC designed and completed clinical and bronchoscopic investigations of patients and volunteers in the Celtic Fire Study; SD co-ordinated clinical data and sample collection and NK managed isolate sequencing; JP designed and completed the time-series analysis of gene expression and metabolite production during airway epithelial differentiation, with bioinformatic analysis from SP and SWO; ET performed microbial community analyses from the Busselton Survey, with contributions from JI-H, LC and MJC; the Survey itself was led by AWM JH, MH, and AJ. WOCM co-ordinated the first draft of the paper, but all authors contributed to the writing and revision.

## Data availability

### Sequences

Raw sequence data for the bacterial isolates have been deposited in the European Nucleotide Archive at the European Bioinformatics Institute under accession number ERP110629. The raw OTU data for the Celtic Fire study is available with the accession number PRJEB40753, and that from the Busselton study with the accession number PRJEB29091.

### Analysis scripts

All data analysis scripts are available online at https://github.com/lcuthber/CelticFire

