## Supplemental Figures for "Genomic and ecologic characteristics of the airway microbial-mucosal complex"

Supplementary Figures

### Supplementary Figure 1. Study Design.

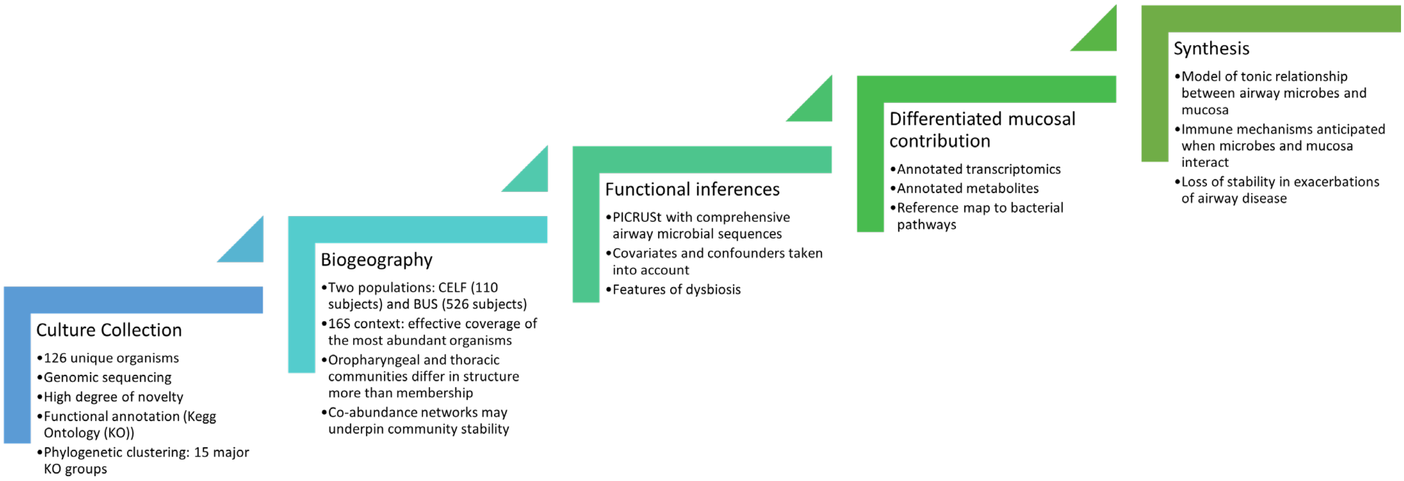

The figure reads from left to right, ascending.

#### SUPPLEMENTARY FIGURE 2. DISTRIBUTION OF NOVEL *STREPTOCOCCUS* SPP. ISOLATES IN GLOBAL *STREPTOCOCCUS* PHYLOGENY.

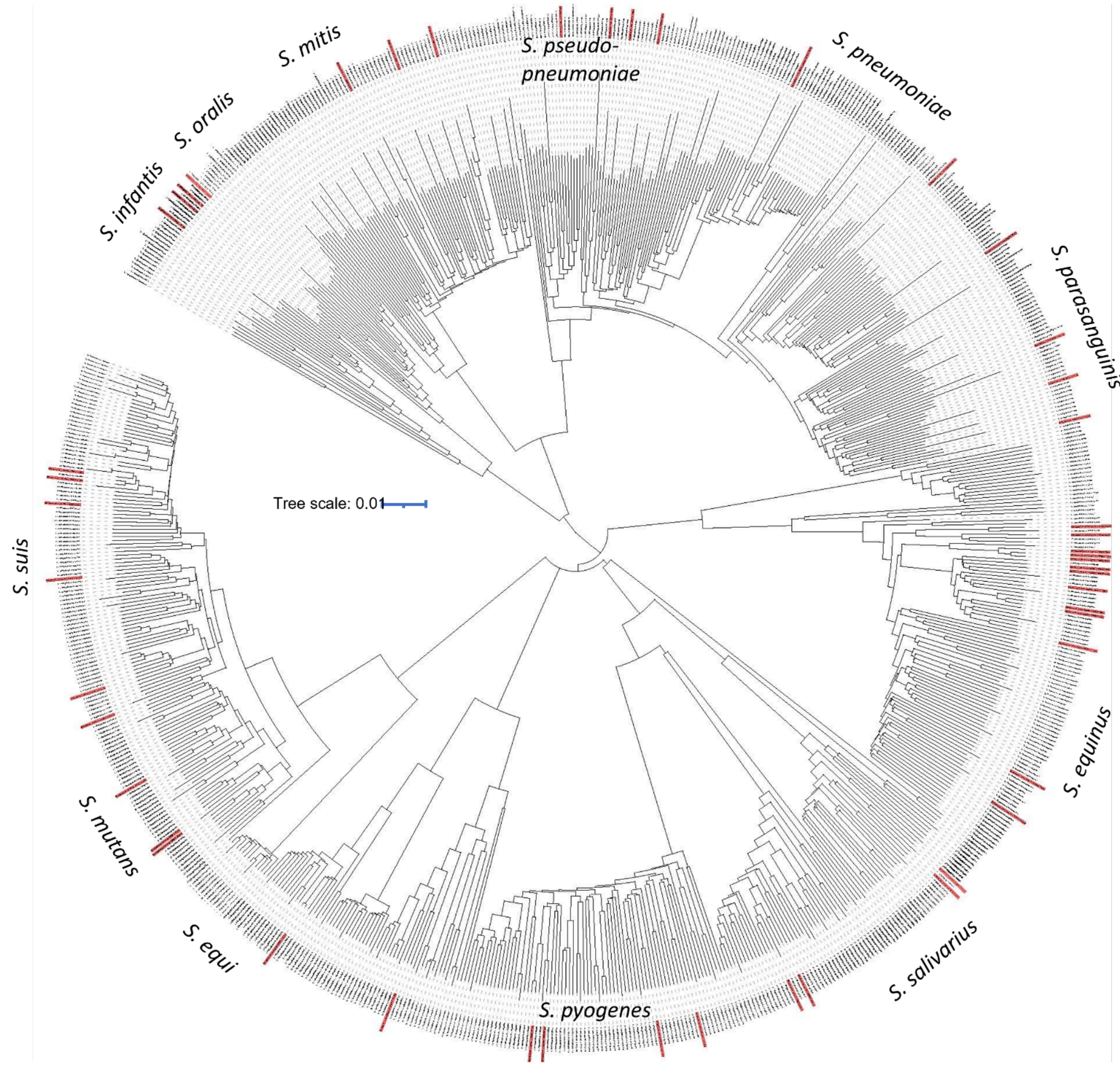

Comparison of the full sequences of airway streptococcal isolates from this study (shown in red) in a pan-genome analysis of 2477 public *Streptococcus* spp. genomes. Airway isolates are each distinct and are widely distributed within the pan-genomes Areas within the tree are labelled according to their most common members, although designations of *S. unknown* are found throughout the tree. A high resolution image is available from Dropbox

### SUPPLEMENTARY FIGURE 3. NETWORK STRUCTURES

#### 3A. CORRELATION STRUCTURE OF BUSSELTON PTOP SAMPLES

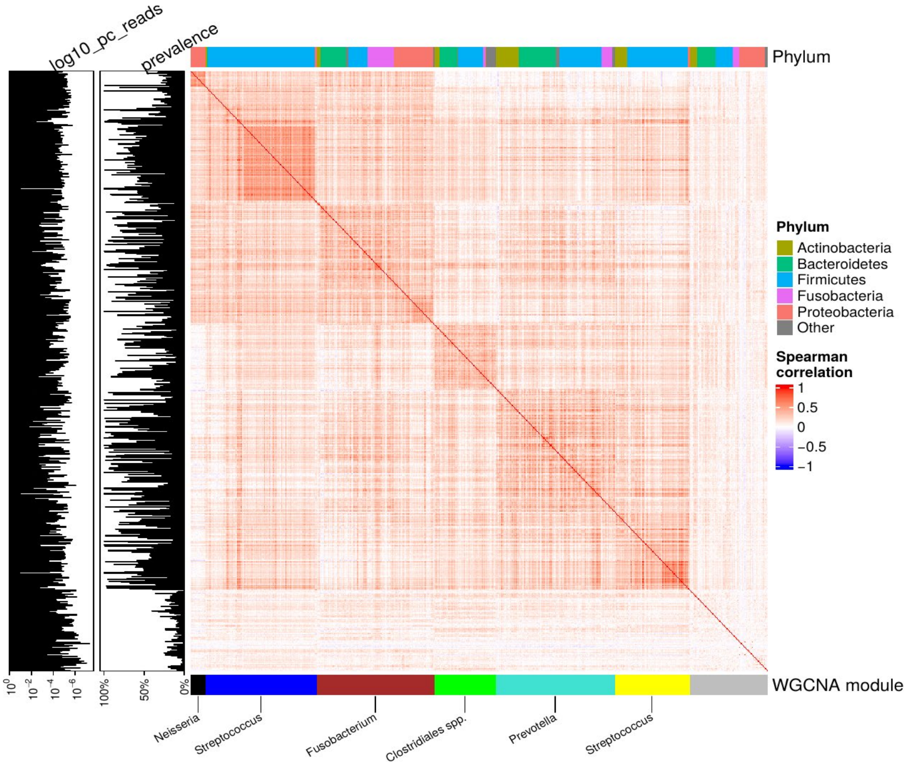

Spearman correlation for the Busselton ptOP samples. The 646 OTUs included in the WGCNA analysis are shown with their Phylum (top colour bar) and WGCNA module (bottom colour bar).

#### 3B. CORRELATION STRUCTURE OF CELF LLL SAMPLES

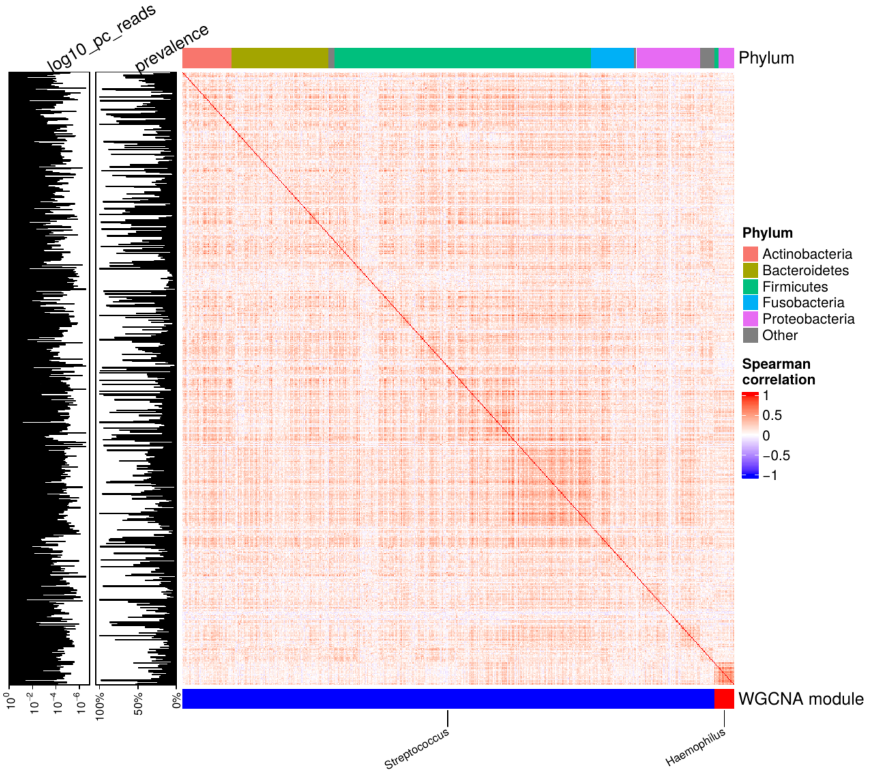

Spearman correlation for the CELF LLL samples. The 646 OTUs included in the WGCNA analysis are shown with their Phylum (top colour bar) and WGCNA module (bottom colour bar). In general, correlations are weaker than in the CELF ptOP (Figure 2d, main paper) and the BUS ptOP samples (3a above).

#### 3C. Conservation of WGCNA modules between sites and studies

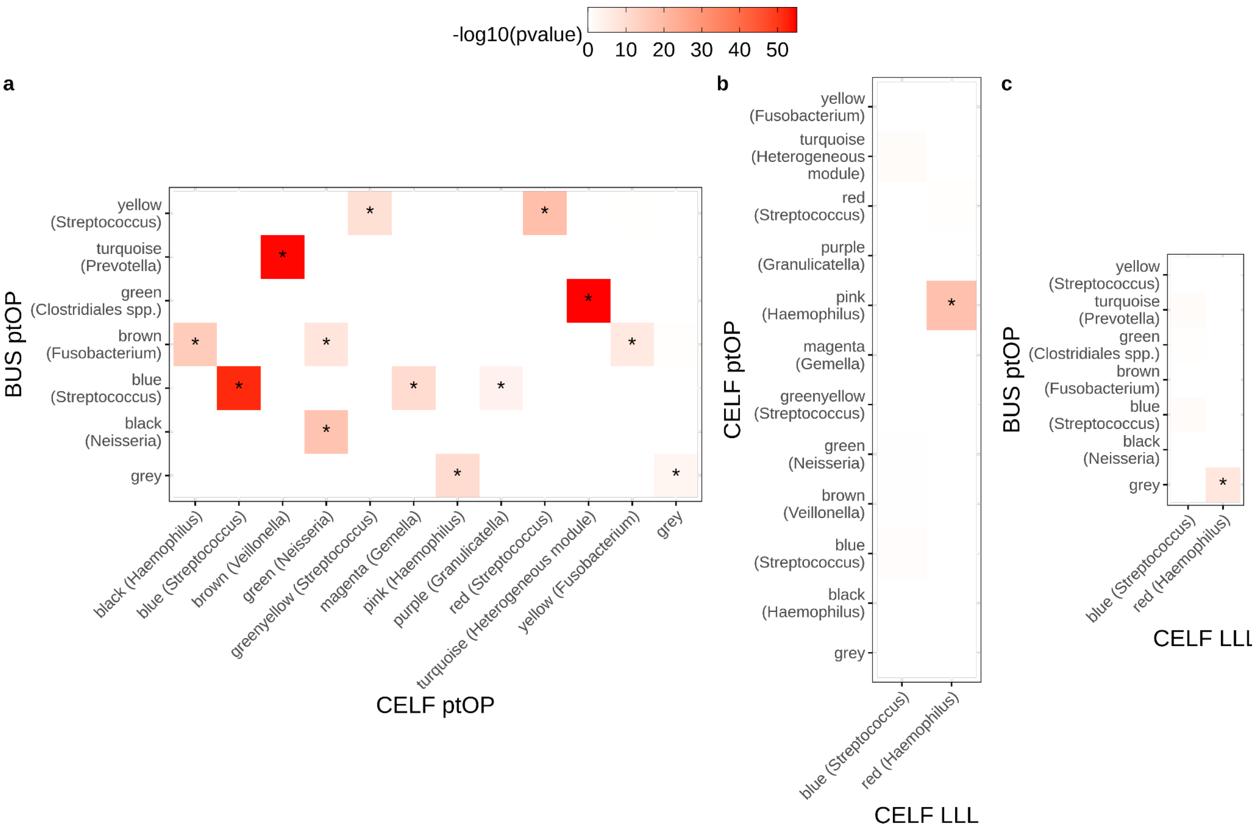

WGCNA network structure is more conserved between the two sets of ptOP samples from the different hemispheres (panel a) than within the same study between the lower airway (panel b) or between the BUS ptOP samples and CELF LLL samples (panel c). Every module in the CELF ptOP network is observed in the BUS ptOP network (p<0.05 denoted using an asterisk). In contrast, only one of the CELF ptOP modules is conserved in the CELF LLL network. Overlap between module assignments tested using Fisher's exact test and corrected using false discovery rate.

#### 3D. Correspondence between *Streptococcus* co-abundance patterns and phylogenetic similarity

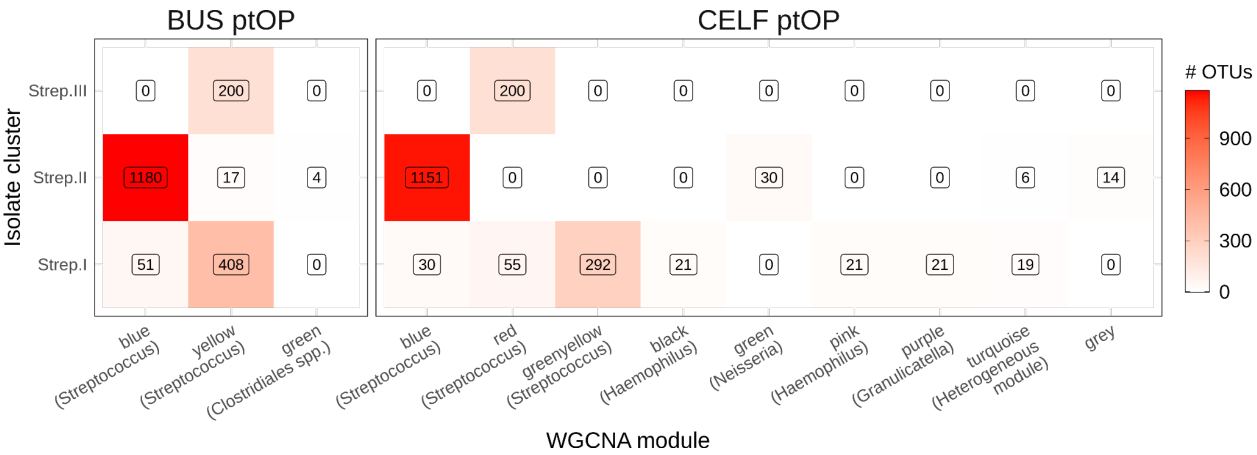

The co-abundance patterns of *Streptococcus* in the ptOP samples mirrors the phylogenetic similarity according to the KO analysis. Plotted is the number of *Streptococcus* OTUs with at least 99% 16S rRNA gene sequence similarity to an isolate, split into the three phylogenetic clusters. In the BUS ptOP samples the blue WGCNA module contains OTUs with higher similarity to isolates in the Strep. II cluster, while the yellow WGCNA module contains OTUs with higher similarity to isolates in Strep. I and Strep. III. In the CELF ptOP samples the blue WGCNA module corresponds to Strep. II, red to Strep. III and greenyellow to Strep. I. In the CELF ptOP network small numbers of *Streptococcus* OTUs are found in other modules.

### SUPPLEMENTARY FIGURE 4. BACTERIAL ISOLATE GENES AND METABOLIC PATHWAYS

#### 4A BACTERIAL GENES MAPPED TO HUMAN METABOLIC PATHWAYS

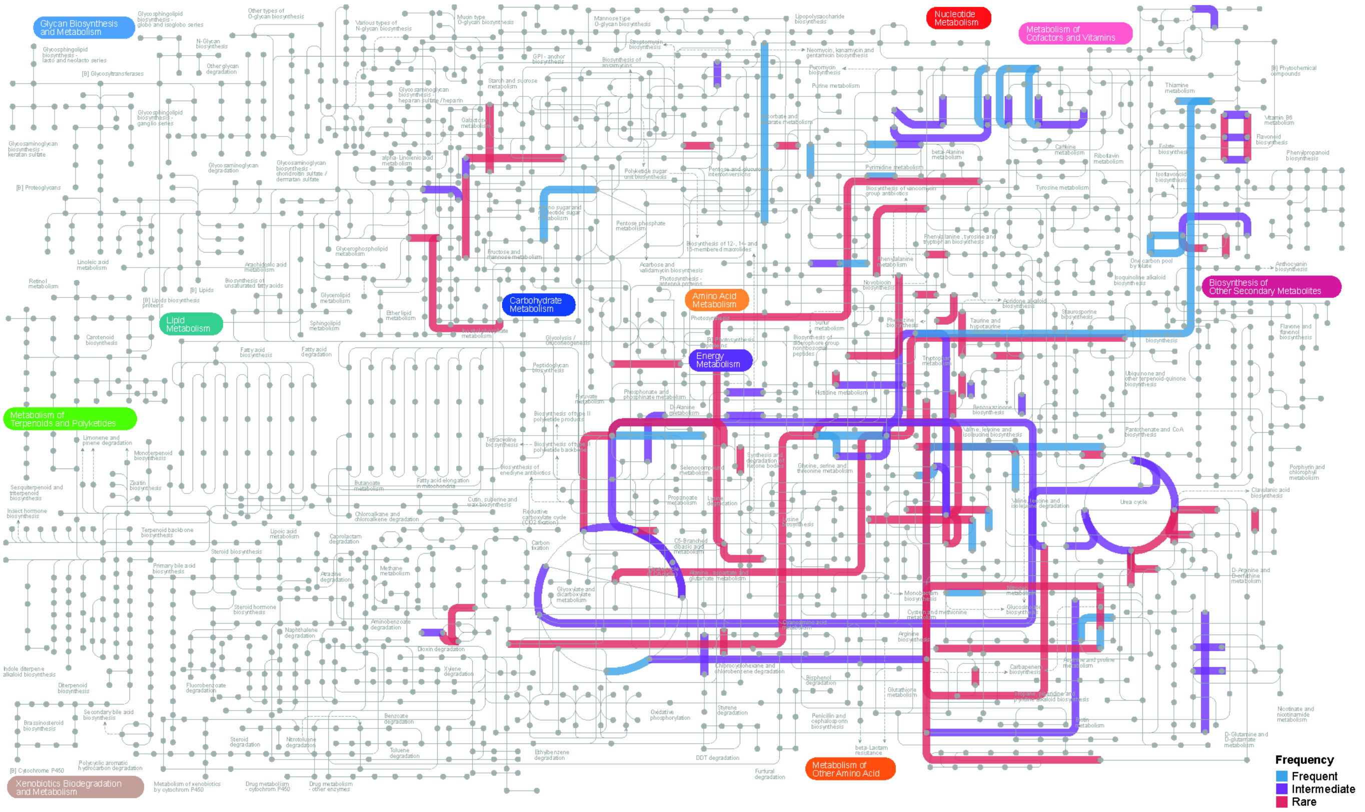

Isolate genes were mapped to human KEGG pathways with the frequency of the genes indicated: ‘frequent’ for genes in >75% of isolates (blue), ‘intermediate’ for genes in 25-75% of isolates (purple) and ‘rare’ for those in <25% of isolates (red). The Plot was produced using iPath 3.0.

### 4B Genes present in bacterial isolates mapped to microbial KEGG pathways

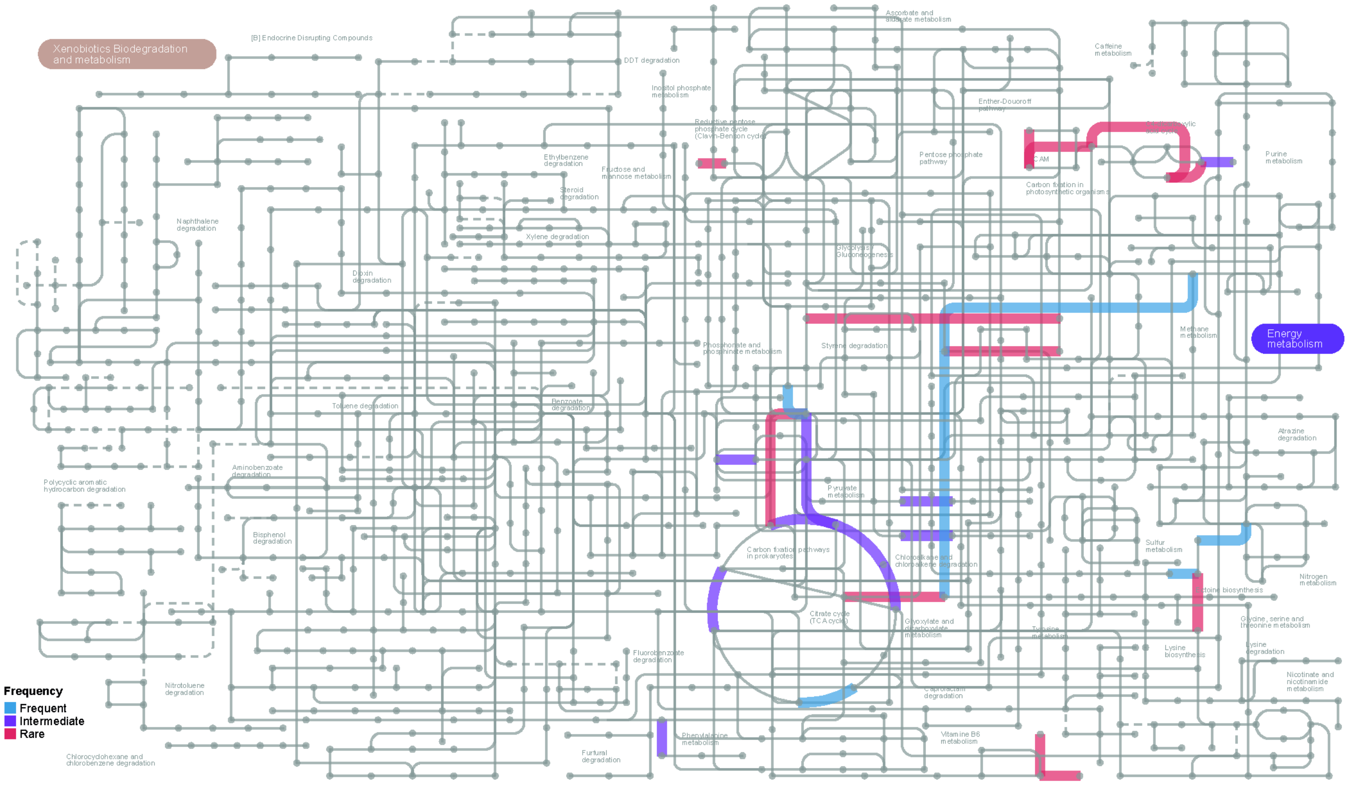

Isolate genes were mapped to microbial KEGG pathways with the frequency of the genes indicated: ‘frequent’ for genes in >75% of isolates (blue), ‘intermediate’ for genes in 25-75% of isolates (purple) and ‘rare’ for those in <25% of isolates (red). The Plot was produced using iPath 3.0.
